# The TBX20-TLE Interaction is Essential for the Second Heart Field

**DOI:** 10.1101/2022.09.27.509637

**Authors:** Whitney Edwards, Olivia K. Bussey, Frank L. Conlon

## Abstract

Tbx20 plays a multifaceted role in cardiac morphogenesis and controls a broad gene regulatory network. However, the mechanism by which Tbx20 activates and represses target genes in a tissue-specific and temporal manner remains unclear. Studies show that Tbx20 directly interacts with the Transducin-like Enhancer of Split (TLE) family of proteins to mediate transcriptional repression of downstream target genes. However, a functional role for the Tbx20-TLE transcriptional repression complex during heart development is not established. To this end, we generated a mouse model with a two-amino acid substitution in the Tbx20 EH1 domain, thereby disrupting the Tbx20-TLE interaction (Tbx20^EH1mut^). We demonstrate that disruption of this interaction impairs critical morphogenic events, including cardiac looping and chamber formation, and ultimately leads to embryonic lethality. Transcriptional profiling of Tbx20^EH1mut^ hearts and analysis of putative Tbx20 direct targets reveals misexpression of the retinoic acid pathway and cardiac progenitor genes, demonstrating that the Tbx20-TLE interaction serves to inhibit cardiac progenitor programs in the developing heart. We find that loss of this interaction also results in perturbations of the second heart field progenitor population, implying that altered cardiac progenitor function may underly the observed cardiac defects in our model. Our studies indicate that TLE-mediated repression is a primary mechanism by which Tbx20 systematically controls gene expression.

## INTRODUCTION

Embryonic heart development requires intricate regulation of transcription factor (TF) regulatory networks (TRNs) that coordinate cardiac cell specification, maturation, and progression of critical morphological events. The essential requirement for cardiac TRNs is emphasized by the observation that mutations in a multitude of cardiac transcription factors are causative in congenital heart disease (CHD), the most common congenital malformation. Clinical and genetic studies provide direct evidence that mutations in TBX20, a T-box TF, are associated with a wide range of cardiac abnormalities. Loss of function mutations in TBX20 are associated with atrial septal defects or mitral valve disease, while gain of function mutations have been reported in patients with Tetralogy of Fallot (i.e., pulmonary outflow tract obstruction, ventricular septal defect, and overriding aortic root)^1–4^.

During embryogenesis, Tbx20 is expressed in several cardiac cell lineages, including the first and second heart field (SHF) cardiac progenitors, endocardial cells, and cardiomyocytes^5,6^. The observed expression of Tbx20 in all major cardiac lineages supports the findings that Tbx20 is involved in numerous developmental processes during embryonic heart development. Global loss of Tbx20 in zebrafish, Xenopus, and mice results in similar phenotypes in which cardiac looping is impaired, and hearts fail to undergo chamber formation, ultimately leading to embryonic lethality^7–12^. In addition, cardiac lineage-specific knockouts reveal that Tbx20 is required for the proliferation and maturation of both the myocardium and endocardial-derived primordial valves (cushions)^13–15^. Together clinical observations and animal model studies highlight an essential and evolutionarily conserved role for Tbx20 in cardiac development and function. However, the mechanisms by which Tbx20 orchestrates these diverse developmental processes during cardiogenesis remains enigmatic.

Identifying the molecular mechanisms by whichTbx20 regulates heart development is complicated by the observation that Tbx20 can act as both a transcriptional activator and repressor. Previous studies suggest that Tbx20 interacts with a network of proteins complexes that dictate its transcriptional activity in a temporal and context dependent manner. A limited number of in vitro studies have shown that Tbx20 interacts with a network of cardiac TFs such as Tbx5, Nkx2.5, Gata4, and Casz1, to synergistically regulate cardiac gene expression^8,16,17^. However, very few interactions have been confirmed and characterized in vivo. Furthermore, the complete network of proteins Tbx20 interacts with during embryonic heart development remains elusive.

Kaltenbrun et al. used an unbiased proteomics-based approach to identify a comprehensive Tbx20 interactome. These studies demonstrated that Tbx20 may mediate transcriptional repression of downstream target genes via its interaction with transducin-like enhancer of split (TLE) proteins, a family of transcriptional co-repressors^18^. The TLE family members are the vertebrate orthologs of the Drosophila Groucho (Gro) protein. TLE/Gro family members are known to play an essential role in diverse developmental processes through their interaction with a myriad of transcription factor families (i.e., Hes, Runx, Nkx, and Fox)^19–23^. TLE/Gro proteins are proposed to mediate transcriptional repression by multiple mechanisms including the recruitment of chromatin remodeling proteins such as histone deacetylases (HDACs) to target gene loci^18,24^. Tbx20 was shown to directly interact with TLE1/3 via an evolutionary conserved N-terminal engrailed homology (EH1) binding motif. In addition, the Tbx20-TLE interaction mediated the recruitment of chromatin remodeling proteins including several members of the nucleosome remodeling and deacetylase complex (NuRD) (Mta1, Rbbp4, Rbbp7 and Hdac2). Further, this study demonstrated that the Tbx20-TLE complex was required for transcriptional repression during Xenopus embryogenesis^18^. These studies provide evidence that Tbx20-TLE interaction mediates transcriptional repression, however, the biological role and requirement for this interaction in cardiac development has not been elucidated.

To investigate the role of the Tbx20-TLE interaction during cardiogenesis we generated a novel mouse model in which a two amino acid substitution was introduced into the Tbx20 EH1 domain, thereby disrupting the Tbx20-TLE interaction (Tbx20^EH1Mut^). Using this model, we demonstrate that the Tbx20-TLE interaction is essential for embryonic heart morphogenesis as mutant mice display impaired cardiac looping and chamber formation ultimately resulting in embryonic lethality. Our transcriptional profiling of Tbx20^EH1Mut^ hearts and analysis of Tbx20 direct targets revealed misexpression of the retinoic acid pathway and progenitor genes, implying that the Tbx20-TLE interaction serves to inhibit cardiac progenitor programs in the developing heart. We report that in the absence of the Tbx20-TLE interaction, there is a reduction of cells in the SHF, and SHF-derived cardiomyocytes fail to differentiate. Collectively these studies define a role for the Tbx20-TLE interaction in the developing heart and suggest that this TLE-mediated repressive program is a primary mechanism by which Tbx20 systematically regulates heart development.

## RESULTS

### The Tbx20-TLE complex is required for cardiac development

Previous studies from our lab demonstrate that a two amino acid substitution (phenylalanine 18 and serine 19 to leucine and isoleucine, respectively) is sufficient to specifically disrupt the Tbx20-TLE interaction and impairs recruitment of chromatin remodeling proteins^18^. To determine the in vivo relevance of the Tbx20-TLE interaction in the developing heart, we used CRISPR/CAS9 genome editing to introduce the same mutation in the mouse germline (Figure 1A). Mice heterozygous for the EH1 amino acid substitutions (Tbx20^+/EH1mut^) are viable, fertile, and phenotypically indistinguishable from control littermates. In contrast, we failed to recover homozygous (Tbx20^EH1Mut^) mice postnatally. (Supplemental Figure 1C). Immunohistochemical analysis showed Tbx20 is expressed in the heart and localized to the nucleus in wild-type and Tbx20^EH1mut^ embryos (Supplemental Figures 1A and 1B). Together, these data indicate that the Tbx20-TLE interaction is essential for embryogenesis.

**Figure 1:**
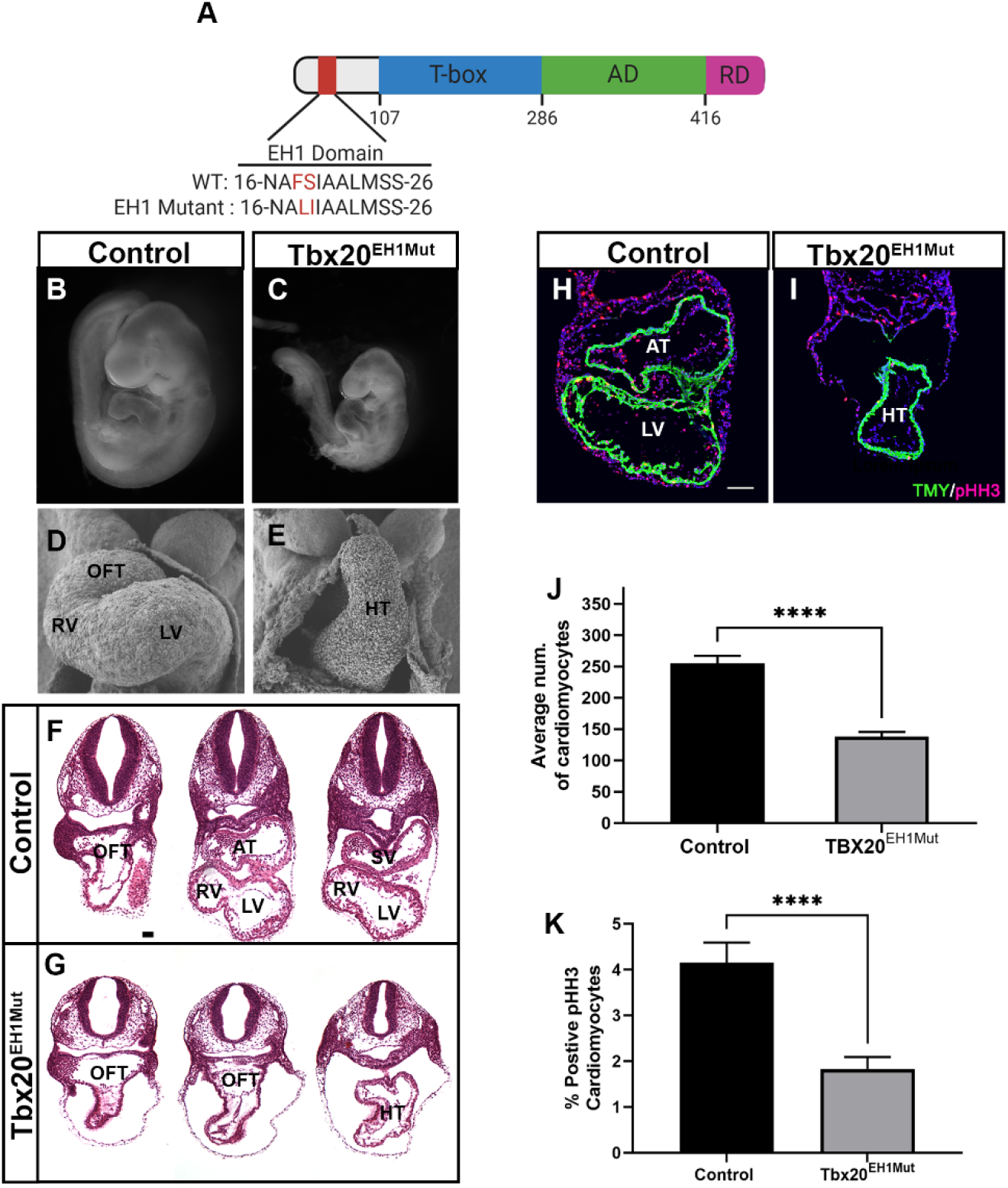
The Tbx20-TLE complex is required for cardiac development. (A) Schematic representation of Tbx20 protein, showing the wild-type and mutant engrailed homology (EH1), T-box, activation (AD), and repression (RD) domains. (B and C) Whole embryo imaging at embryonic day 9.5 (E9.5) shows growth retardation and altered cardiac morphology in Tbx20^EH1Mut^ embryos compared to control. N≥3 per genotype. (D and E) Scanning electron microscopy analysis at E9.5 reveals impaired cardiac looping and chamber formation of Tbx20^EH1Mut^ hearts compared to control. 400x magnification. N=2 per genotype. (F and G) Hematoxylin and Eosin stained transverse sections representing the anterior, middle and posterior regions of E9.5 control and Tbx20^EH1Mut^ hearts. Scale bar, 66 μM. N=3 per genotype. (H and I) Immunohistochemical analysis of phosphohistone-H3 (pHH3) positive cardiomyocytes (Tropomyosin; TMY) reveals a significant decrease in the cardiomyocyte mitotic index in Tbx20^EH1Mut^ hearts compared to control. Scale bar, 100 μM. (J) Quantitation of average number of cardiomyocytes in control and Tbx20^EH1Mut^ hearts at E9.5. N=3-4, per genotype. (K)Quantitation of cardiomyocytes mitotic index in control and Tbx20^EH1Mut^ hearts at E9.5. N=3-4, per genotype. Data are expressed as mean values, error bars represent ± standard error of the mean (SEM). ****p≤ 0.0001. OFT, outflow tract; RV, right ventricle; LV, left ventricle; AT, atria; SV, sinus venous; HT, heart tube.

Gross morphological analysis of Tbx20^EH1Mut^ embryos at embryonic day 9.5 (E9.5) revealed mutant embryos displayed pericardial edema and hemorrhaging (Figure 1B and 1C). Ultrastructural imaging analysis (scanning electron microscopy) showed Tbx20^EH1Mut^ hearts initiate heart tube formation but failed to undergo cardiac looping or cardiac chamber formation (Figure 1D and 1E). Further, in contrast to wild-type embryos, Tbx20^EH1Mut^ hearts fail to form identifiable cardiac regions or chambers including the outflow tract (OFT), left and right ventricles, and atria (Figures 1F and 1G). Moreover, the Tbx20^EH1mut^ cardiac phenotype was accompanied by a significant decrease in cardiomyocyte number (Tropomyosin+;TMY+) and a concomitant decrease in the mitotic index (pHH3+/TMY+) (Figure 1H and 1K). Taken together, this data reveals that the Tbx20-TLE interaction is required for heart formation and viability.

### TBX20-TLE interaction functions to repress retinoic acid signaling

TLE proteins function as a transcriptional co-repressors, therefore we hypothesize that the Tbx20-TLE interaction functions to repress inappropriate gene expression in the developing heart^20,22–24^. To test this hypothesis, we conducted transcriptional profiling (RNA-Sequencing) on wild-type and Tbx20^EH1mut^ hearts at E9.5. Our analysis identified 2,218 differentially expressed genes (DEGs) (adjusted p-value ≤ 0.05 and log2 fold change ≥ ±1) of which, 1,363 genes were significantly up regulated, and 855 genes were down regulated in Tbx20^EH1Mut^ hearts compared to controls (Figure 2A). To assess the molecular pathways associated with DEGs, Ingenuity Pathway Analysis (IPA) was performed (Figure 2B). Pathways associated with down regulated genes included “cardiac hypertrophy signaling” and “factors promoting cardiogenesis in vertebrates” all of which contain genes involved in cardiomyocyte function and growth (*Myh6, Nppa, Scn5a, Gja5, Hand1, Bmp2)*. In addition, the “cyclins and cell cycle regulation” pathway included several genes involved in the regulation of cell cycle progression and proliferation (*ccnb1, ccnd1, ccnd2*). Down regulation of these pathways corroborates our findings that cardiomyocyte number and mitotic index were decreased in Tbx20^EH1Mut^ hearts.

**Figure 2:**
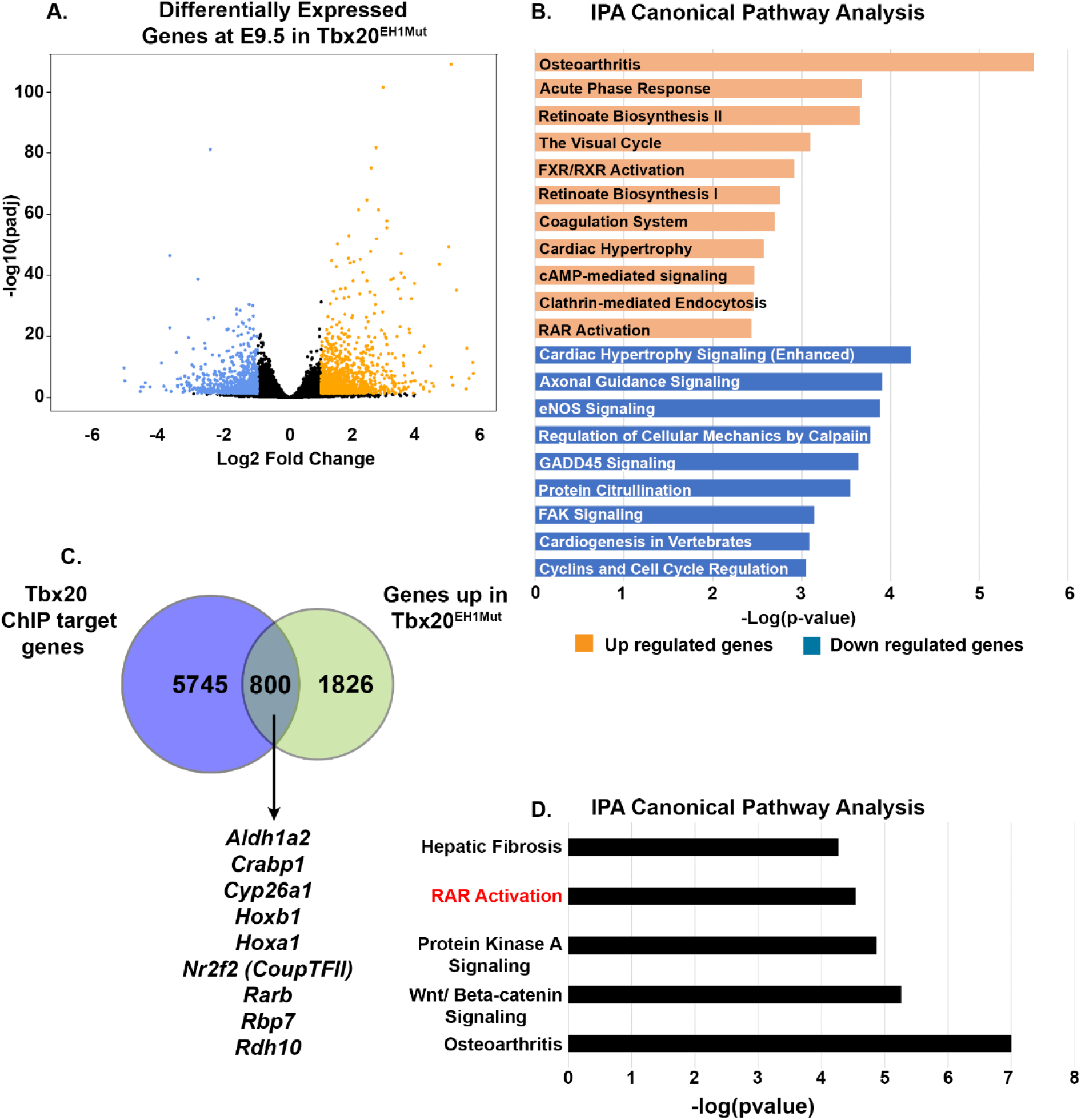
TBX20-TLE interaction functions to repress retinoic signaling. (A) Volcano plot of genes identified by RNA-sequencing to be differentially expressed between WT and Tbx20^EH1Mut^ hearts at E9.5 (adjusted p-value ≤ 0.05 and log2 fold change ≥ ±1). Down regulated genes are shown in blue and up regulated genes shown in orange. (B) Ingenuity pathway analysis (IPA) of differentially expressed genes reveals misexpression of retinoic acid associated genes in Tbx20^EH1Mut^ hearts compared to controls. (C) Overlap of up regulated genes in Tbx20^EH1Mut^ hearts (adjusted p-value ≤ 0.05 and log2 fold change ≥ 0.5) with Tbx20 ChIP-seq (chromatin immunoprecipitation followed by high throughput sequencing) dataset^13^. (D) Subset of genes up regulated in Tbx20^EH1Mut^ hearts and identified as putative direct Tbx20 targets by Chip-Seq are associated with retinoic acid signaling by IPA.

We hypothesize that Tbx20 recruits TLE family members to target genes to mediate transcriptional repression during embryonic heart development, therefore we focused on genes and pathways that were aberrantly up regulated in Tbx20^EH1Mut^ hearts. Surprisingly, analysis of up regulated genes identified several pathways associated with retinoic acid (RA) signaling including “Retinoate Biosynthesis I”, “Retinoate Biosynthesis II”, and “RAR Activation”. Genes within these biological categories included the RA-synthesizing enzyme (*Aldh1a2)*, several RA binding proteins (*Rpb1, Rbp2, Rbp7)*, and retinol dehydrogenase enzymes (*Rdh12)*.

The RA signaling pathway is essential for cardiac morphogenesis and perturbations in the RA pathway are associated with CHD including malformations of the outflow tract (OFT), septal defects and cardiac looping defects^25–31^. Therefore, we sought to determine if components of the RA signaling pathway are direct transcriptional targets of Tbx20. To this end, we overlaid our RNA-Seq data (adjusted p-value ≤ 0.05 and log2 fold change ≥ 0.5) with previously published Tbx20 ChIP-Seq (chromatin immunoprecipitation with high throughput sequencing) generated from embryonic hearts. We identified 800 genes that were putatively bound by Tbx20 and up regulated in the Tbx20^EH1Mut^ hearts (Figure 2C). IPA analysis of these putative direct targets once again revealed genes associated with RA signaling (Figure 2D). Most notably, *Aldh1a2* was identified as a putative direct Tbx20 target. In aggregate, our data suggests that the Tbx20-TLE complex suppresses aberrant RA signaling during embryonic heart development.

### Cardiac progenitors are arrested in development in Tbx20^EH1mut^ hearts

Misexpression of RA pathway genes in Tbx20^EH1Mut^ hearts was unexpected as these genes are primarily expressed in second heart field (SHF) cardiac progenitors located outside of the primary heart tube at E9.5. This finding in combination with impaired looping and chamber formation in Tbx20^EH1Mut^ hearts may indicate arrested cardiomyocyte development.

To test this possibility, we overlapped our gene expression dataset with a subset of genes shown to be specifically enriched in multipotent cardiac progenitors (MP) as identified by single-cell RNA-sequencing^32^. We find that approximately 64% of MP enriched genes (25/39) display differential gene expression, all of which are overexpressed in the Tbx20^EH1mut^ heart tube (Figure 3A). These included *Isl1, Fgf8*, and *Osr1*, well established markers of the cardiac progenitor population (Figure 3B)^28,33,42,43,34–41^. Collectively, these data imply that the Tbx20-TLE interaction functions to repress expression of RA signaling genes within the developing heart tube, thereby promoting proper timing of cardiomyocyte differentiation.

**Figure 3:**
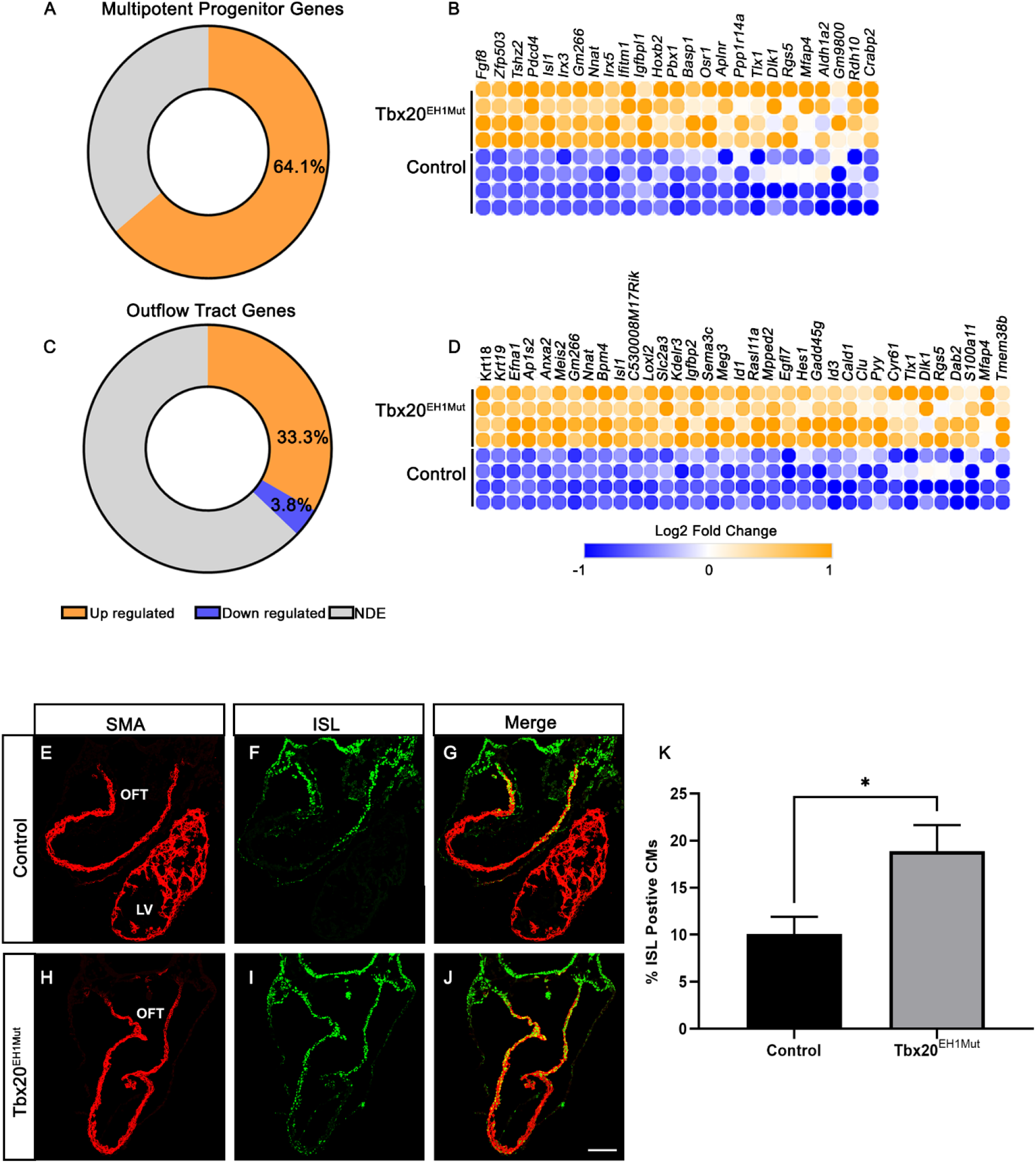
Cardiac progenitors are arrested in development in Tbx20^EH1mut^ hearts. (A) Percent of differentially expressed multipotent progenitor-associated genes in Tbx20^EH1Mut^ hearts. (B) Heatmap comparing expression of multipotent progenitor-associated genes in Tbx20^EH1Mut^ and control hearts. (C) Percent of differentially expressed outflow tract-associated genes in Tbx20^EH1Mut^ hearts. (D) Heatmap comparing expression of outflow tract-associated genes in Tbx20^EH1Mut^ and control hearts. (E-J) Immunohistochemical analysis shows significant increase in the percent of Islet1 (Isl1; green) positive cardiomyocytes (smooth muscle actin; SMA; red) in Tbx20^EH1Mut^ hearts compared to controls. Scale bar, 100 μM. (K) Quantitation of Isl1-positive cardiomyocytes in control and Tbx20^EH1Mut^ hearts at E9.5. N=3, per genotype. Data are expressed as mean values, error bars represent ± standard error of the mean (SEM). *p≤ 0.05. OFT, outflow tract; LV, left ventricle; NDE, Not differentially expressed.

### The Tbx20-TLE complex regulates second heart field derived cells

During cardiac development SHF progenitors migrate to the anterior and posterior poles of the developing heart tube and give rise to, the OFT, right ventricle, and the arterial poles the heart^27^. The RA signaling pathway is known to regulate development of OFT. Our findings that this pathway is up regulated in Tbx20^EH1mut^ hearts, leads us to query if the Tbx20-TLE interaction is essential for the development of the OFT. Therefore, we overlapped our gene expression dataset with genes that are specifically enriched in the OFT during embryonic heart development. Our analyses revealed approximately 33% (35/105) of OFT enriched genes were upregulated in Tbx20^EH1mut^ hearts versus controls (Figure 3C). Up regulated genes include *Isl1, Sema3c, Meis2*, and *Hes1*, critical mediators of OFT development (Figure 3D)^33,44,45^.

Our analysis of MP-and OFT-enriched genes identified up regulation of *Isl1* in Tbx20^EH1Mut^ hearts. Isl1, a marker of both the SHF progenitors and the OFT, is a master regulator of cardiac progenitor proliferation, migration, and survival, and was previously identified as a direct target of Tbx20^33,36^. Isl1 is highly expressed in SHF progenitors, its expression is maintained as myocardial progenitors integrate into the forming heart tube and is eventually downregulated as cells begin to differentiate. To assess and validate overexpression of Isl1 we performed immunohistochemical analysis of WT and Tbx20^EH1Mut^ hearts at E9.5. In WT hearts Isl1-positive cardiomyocytes (Isl1+SMA+) are primarily located in the proximal OFT (Figure 3E-3G). In contrast, the number of Isl1+SMA+ cells are expanded throughout the heart tube in Tbx20^EH1Mut^ hearts (Figure 3H-3J). Furthermore, we observed an almost 2-fold increase in the percent of Isl1+SMA+ cardiomyocytes in Tbx20^EH1Mut^ hearts compared to controls (Figure 3K). These findings suggest that SHF-derived cells fail to undergo differentiation and remain in a cardiac progenitor-like state.

The primary heart tube is derived from first heart field (FHF) cells, an additional pool of cardiac progenitors. Our findings that SHF-derived cells are increased in the heart may suggest an imbalance in cells derived from these two populations. To test this hypothesis, we probed our gene expression dataset for FHF-associated genes. Intriguingly, we find hallmark genes of the FHF, including *Hcn4, Gata4*, and *Hand1*, are down regulated in Tbx20^EH1mut^ hearts compared to controls^46–49^. In aggregate, these findings imply that the Tbx20-TLE interaction is essential for the development of first- and second-heart field derived cells.

### TLE family members are expressed in the SHF during embryonic heart development

Studies of SHF development have shown that perturbations of SHF progenitors result in impaired heart tube elongation and cardiac looping defects, similar to those we observe in Tbx20^EH1mut^ mice. Previous studies demonstrate that *Tbx20* is expressed in the SHF progenitors during embryonic heart development. We therefore hypothesized that the Tbx20-TLE interaction regulates the SHF progenitor population. To address this hypothesis, we first examined the expression of TLE family members in the SHF during embryonic heart development.

The SHF can be subdivided into two distinct regions. Progenitors derived from the anterior SHF (aSHF) give rise to right ventricle and portions of the OFT. In contrast, the posterior SHF (pSHF) progenitors primarily give rise to the atria and a subset of cardiac vessels. Analysis of previously published transcriptomic profiling of the aSHF and pSHF at E9.5 revealed that *Tle1* and *Tle3* are highly expressed in both SHF domains (Figure 4A)^50^. *Tbx20* is also detected in both SHF populations however its expression pattern more closely parallels well-established pSHF makers (*Aldh1a2, Tbx5* and *Osr1*) as we observe higher expression in the pSHF compared to the aSHF (Figure 4A). We obtained similar results from a recent single-cell RNA-seq analysis of the *Isl1*-postive SHF progenitor population at early stages of embryonic heart development (E7.75, E8.25, E9.25) (Figure 4B-4E)^32^. In addition, immunohistochemical analysis of wild-type embryos at E9.5 validates the expression of TLE family members in the SHF. Our analysis demonstrates Tle3 is highly expressed in both domains of the SHF and colocalizes with the pan-SHF marker Isl1 in the vast majority of SHF cells. Together, these data demonstrate that *Tbx20*, Tle1, Tle3, and Isl1 are expressed in SHF progenitors.

**Figure 4:**
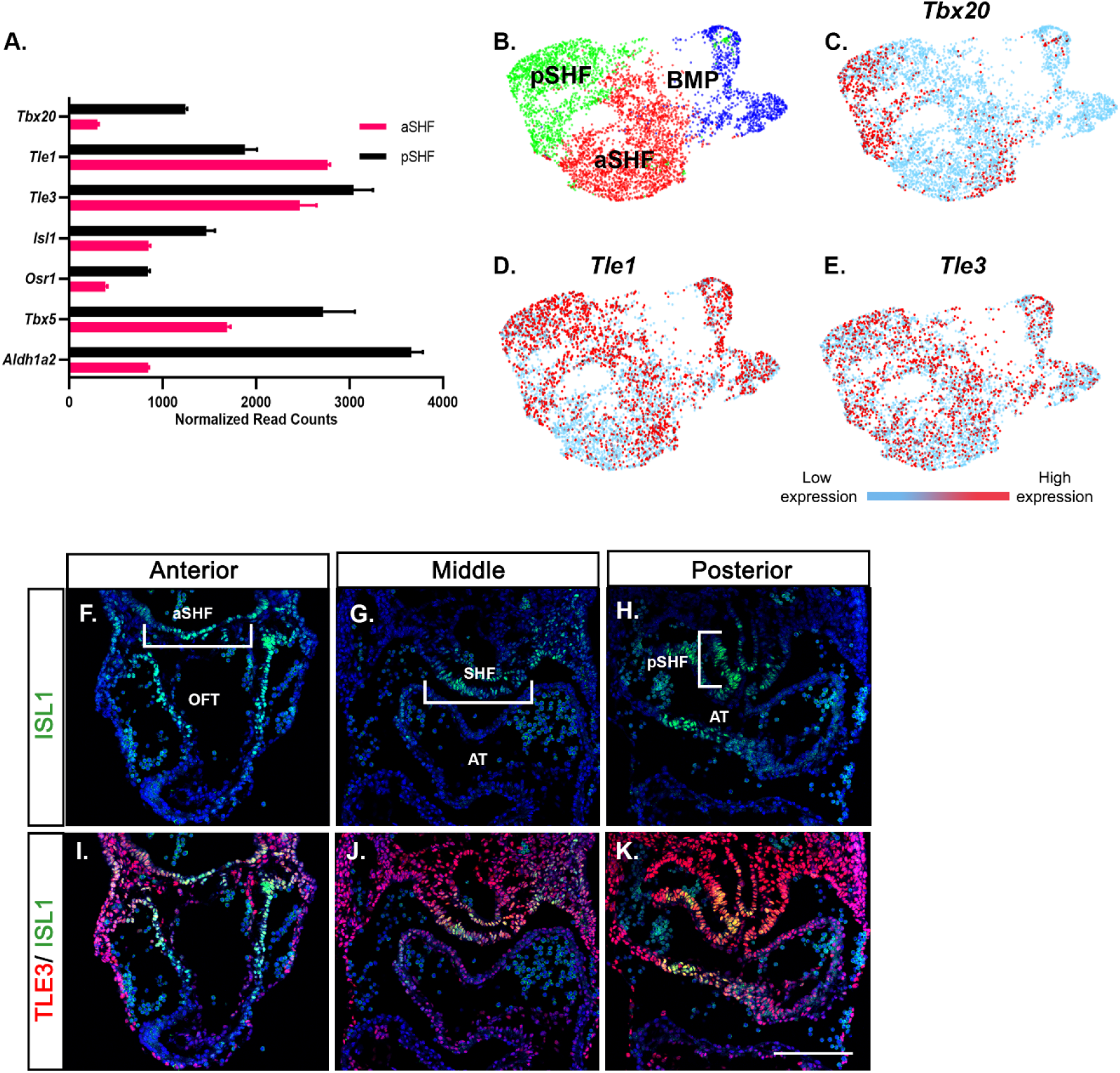
TLE family members are expressed in the SHF during embryonic heart development. (A) Relative transcript abundance of *Tbx20, Tle1, Tle3* and SHF-associated genes from RNA-seq analysis of anterior and posterior SHF progenitor populations^50^. (B) Uniform manifold approximation and project (UMAP) plot of cardiac progenitor subpopulations generated from single-cell RNA-seq analysis of embryonic hearts^32^. (C-E) UMAP plot of *Tbx20, Tle1*, and *Tle3* in cardiac progenitor subpopulations. (F-K) Immunohistochemical co-expression analysis of Isl1 (green) and Tle3 (red) shows Tle3 is robustly expressed in the SHF of control embryos at E9.5. Scale bar, 100 μM. aSHF, anterior second heart field; pSHF, posterior second heart field; BMP, branchiomeric muscle progenitors; OFT, outflow tract; AT, atria.

### Tbx20-TLE is essential for maintenance of the SHF

In combination, our findings that Tbx20^EH1Mut^ embryos display an impaired elongation and looping phenotype, the upregulation of cardiac progenitor and OFT associated genes in the heart tube, and the detection of TLE family member in the SHF suggest the SHF progenitor population may be affected in Tbx20^EH1Mut^ embryos.

To determine if the SHF is altered in E9.5 Tbx20^EH1Mut^ embryos we first quantified the expression of Isl1 which labels both the aSHF and pSHF. Our analysis revealed a significant reduction in the total number of Isl1-postive (Isl1+) cells in the SHF in Tbx20^EH1Mut^ embryos compared to wild-type controls (Figure 5A-5G). In addition, we determined the mitotic index (pHH3+Isl1+) of Isl1+ SHF progenitors was also reduced in Tbx20^EH1Mut^ embryos compared to controls (Figure 5B, 5E, and 5H).

**Figure 5:**
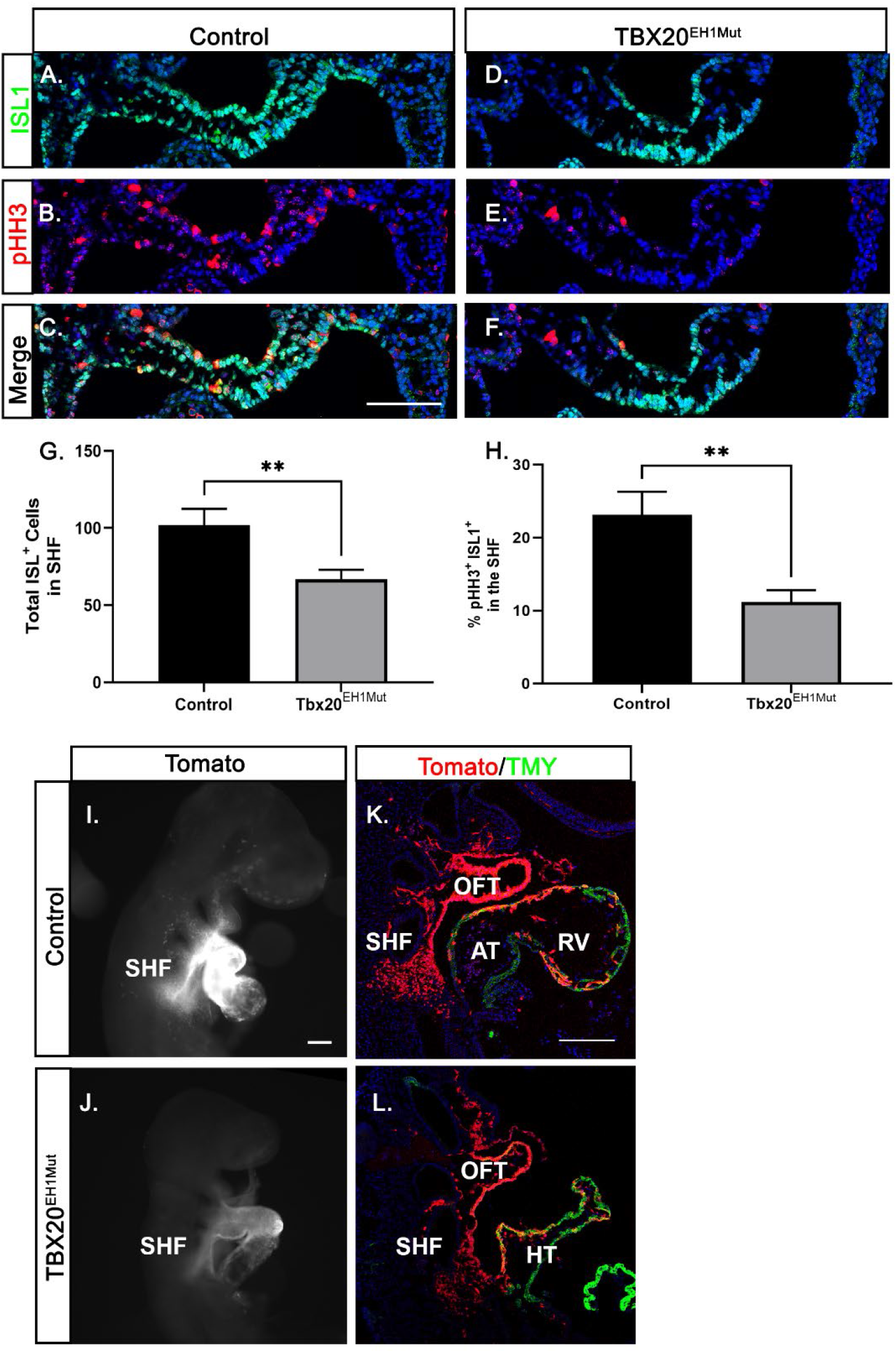
Tbx20-TLE is essential for maintenance of the SHF. (A-F) Immunohistochemical analysis of phosphohistone-H3 (pHH3; red) and Isl1 (green) expression in the SHF shows a significant decrease in the mitotic index of Isl1-positive cells in the SHF of Tbx20^EH1Mut^ embryos compared to controls at E9.5. Scale bar, 100 μM. (G) Quantitation of total number of Isl1 positive cells in the SHF of control and Tbx20^EH1Mut^ embryos. (H) Quantitation of mitotic index of Isl1 positive cells in the SHF of control and Tbx20^EH1Mut^ embryos. (I and J) Whole embryo fluorescent imaging of Mef2c genetic lineage tracing (Tomato fluorescence is pseudo-colored white) shows a reduced SHF progenitor population in Tbx20^EH1Mut^ embryos compared to control. Scale bar, 100 μM. (K and L) Representative sagittal sections of Mef2c genetic lineage tracing (red) in control and Tbx20^EH1Mut^ embryos. Sections are co-stained with TMY (green). Scale bar, 200 μM. Data are expressed as mean values, error bars represent ± standard error of the mean (SEM). **p≤ 0.01. SHF, second heart field; OFT, outflow tract; RV, right ventricle; AT; atria; HT, heart tube.

To directly examine the fate of the SHF derived cells in Tbx20^EH1mu^ embryos, we lineage traced SHF cells in control and Tbx20^EH1mut^ embryos. For these studies, the Tbx20^EH1Mut^ mouse line was crossed to *Mef2c-AHF-Cre*;Rosa^td^ mice, a transgenic mouse line that drives Tomato expression in the a sub-domain of SHF progenitors and lineages derived from this population^51^. Tomato-positive (Td+) cells are primarily detected in the SHF, OFT and RV in control embryos (Figure 5E and 5F). In Tbx20^EH1Mut^ mice, Td+ cells are primarily detected in the heart tube, and label what is presumed to be the OFT. These data demonstrate that at least a subset of SHF progenitors can migrate and integrate into the heart tube in Tbx20^EH1Mut^ embryos. However, we also observed a striking reduction in the SHF Td+ population in Tbx20^EH1Mut^ mice compared to control embryos (Figure 5G and 5H). These data reveal that loss of this critical progenitor population during development likely contributes to the cardiac phenotype observed in Tbx20^EH1Mut^ embryos.

## DISCUSSION

Tbx20 is an essential transcription factor for heart development, and its disease relevance is well established. However, there are many critical questions unanswered about the mechanism by which Tbx20 functions. Here, we have introduced a two amino acid substitution into the Tbx20 EH1 motif, thereby preventing the interaction of Tbx20 with the TLE co-repressors. Our studies demonstrate that the Tbx20-TLE interaction is required for cardiac morphogenesis and is essential for maintenance of the SHF progenitor population.

The mechanism by which Tbx20 systematically regulates transcriptional repression is poorly understood. Our studies indicate that Tbx20 recruitment of TLE proteins may be a primary mechanism by which Tbx20 mediates transcriptional repression. Tbx20 has been shown to directly repress Isl1 in differentiating cardiomyocytes^9,14^. Concordantly, we show that in Tbx20^EH1mut^ hearts the number of Isl1 positive cardiomyocytes is significantly increased. This finding suggests that Tbx20 repression of Isl1 is TLE-mediated. In addition, a large subset of multipotent cardiac progenitor genes, including several RA pathway genes, were shown to be misexpressed in Tbx20^EH1mut^ hearts. Taken together, our data suggests that the Tbx20-TLE interaction is required to repress expression of cardiac progenitor genes. Furthermore, these findings indicate that impaired cardiomyocyte maturation and differentiation likely underly the chamber maturation phenotype observed in Tbx20^EH1mut^ hearts.

SHF progenitors, located dorsomedially to the primary heart tube, migrate to the anterior and posterior poles of the heart, differentiate, and drive tube elongation. Altered expression of SHF genes including *Nkx2*.5, *Isl1*, and *Mef2c* is associated with impaired heart tube elongation and cardiac looping defects^33,52,53^. Previous research supports a role for Tbx20 in cardiac progenitor function and maintenance. In vitro studies demonstrate that Tbx20 functions in concert with other cardiac progenitor transcription factors to regulate expression of SHF-associated genes^11^. In addition, recent studies in zebrafish show that loss of Tbx20 early in cardiogenesis results in impaired cardiac progenitor development^54^. In support of these studies, we find that loss of the Tbx20-TLE interaction results in a marked reduction in the SHF progenitor population.

How and when does the Tbx20-TLE interaction function in the SHF? We favor a model by which the aSHF is decreased due to the mis-patterning of the anterior and posterior SHF. This hypothesis is consistent with studies that have shown an increase in RA signaling in the pSHF leads to a reduction in aSHF (Tbx1, Fgf8, Fgf10 etc.) gene expression^38,55–58^. In Tbx20^EH1mut^ embryos, alterations in RA signaling disrupt SHF patterning and in turn leads to the loss of cardiac progenitors. The remaining SHF progenitors that survive and integrate into the heart tube are mis-programmed and therefore are unable to properly differentiate We note that the phenotype of the Tbx20^EH1mut^ heart is distinct from that reported for Tbx20 nulls^9–12^. Moreover, we have identified Tbx20-TLE targets genes distinct from those observed in Tbx20 null hearts. These findings indicate that Tbx20 acts through additional chromatin remodeling complexes during cardiogenesis. Components of the Nucleosome Remodeling and Deacetylase complex and the chromatin remodeling ATPase RUVBL1/RUVBL2 complex have been shown to interact with Tbx20 and may account for the additional transcriptional functions of Tbx20^18^. However, in vivo studies for the function of these interactions have yet to be reported.

Our studies indicate the Tbx20-TLE interaction is essential for heart development. Therefore, components of the TLE repression complex and the transcriptional targets of the Tbx20-TLE complex, may be causative in a range of congenital heart diseases. Our findings also infer that patient phenotypes due to mutations in the cardiac Tbx20-TLE complex or downstream target genes are not only due to loss of gene expression but also misexpression of genes in SHF derived cells. Finally, it is noted that Tbx20 is not only involved in CHD but has also been shown to be associated with cardiomyopathy in patients^1,17,59^. Consistently, it has been demonstrated in mice that ablation of Tbx20 in adult cardiomyocytes leads to the onset of severe cardiomyopathy and premature death^11,60^. Determining the role of the Tbx20-TLE in adult-onset heart disease and defining mutations in the TLE/Groucho complex will be critical for assessing the role of the Tbx20-TLE interactions in additional disease states.

## ACKNOWLEDGEMENTS

We thank the UNC Animal Models Core for help in the generation of the Tbx20^EH1Mut^ allele. We thank the UNC Microscopy Services Laboratory Core for the processing and imaging of the transmission electron microscopy.

## FIGURE LEGENDS

**Supplemental Figure 1:**
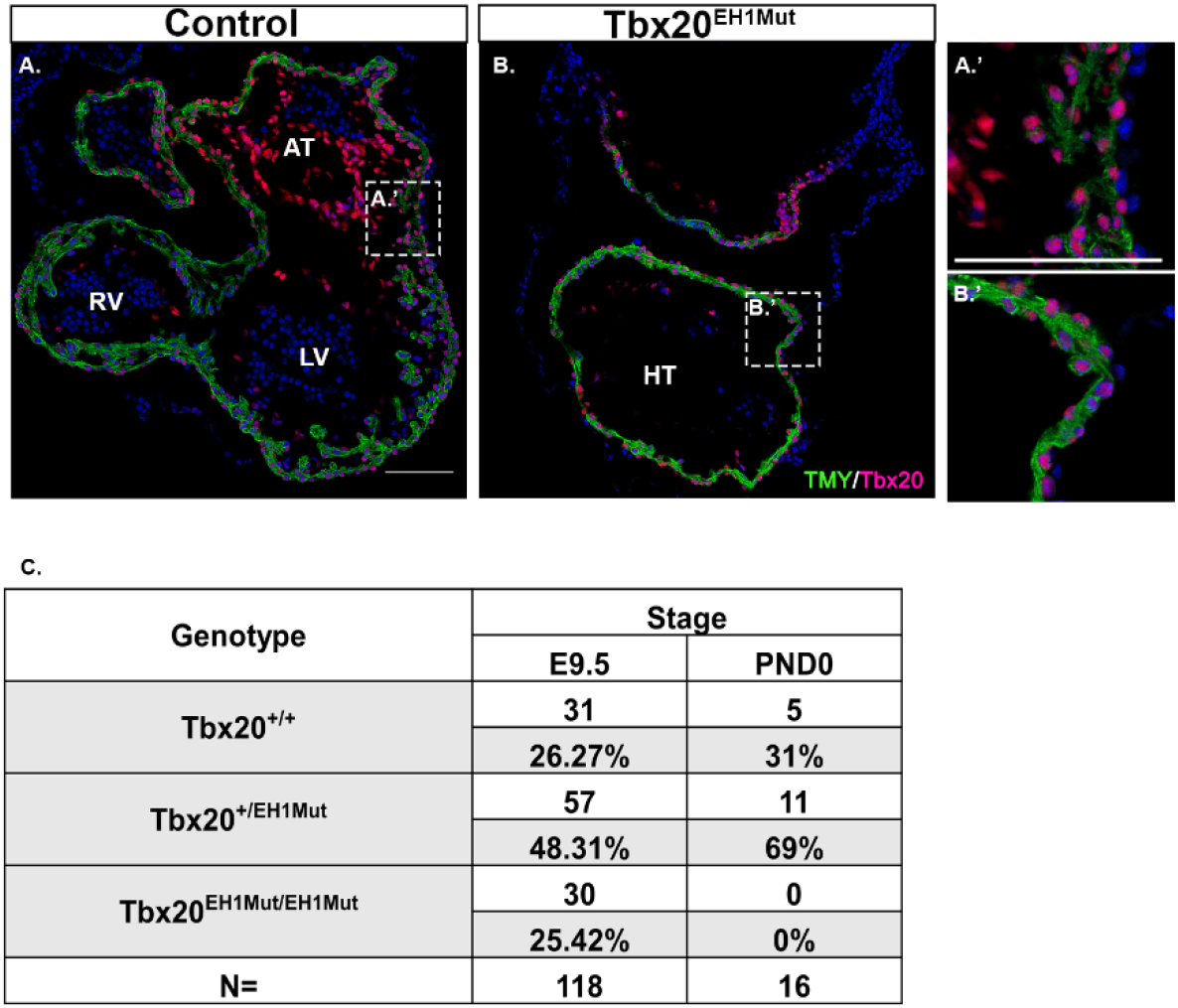
(A-B’) Immunohistochemical analysis of Tbx20 (red) positive cardiomyocytes (TMY; green) in control and Tbx20^EH1Mut^ hearts at E9.5. Scale bar, 100 μM. (C)Genotype distribution of wild-type, heterozygous and homozygous mutant embryos at E9.5 and postnatal day (PND) 0.

## MATERIALS AND METHODS

### Mice

Tbx20^EH1mut^ mice were generated by the UNC Animals Models Core Facility. Wild-type C57/Bl6, Gt(ROSA)26Sortm14(CAG-tdTomato)Hze (stock 007194), Mef2c-AHF-cre^61^ mice have all been previously described and were obtained from The Jackson Laboratory. Research was approved by the Institutional Animal Care and Use Committee at the University of North Carolina and conforms to the Guide for the Care and Use of Laboratory Animals.

### Generation of Tbx20^EH1mut^ mouse line

CRISPR/Cas9 genome editing was used to introduce the following mutation in the Tbx20 engrailed homology domain (EH1): phenylalanine 18 and serine 19 to an leucine and a isoleucine, respectively.

Cas9 guide RNAs flanking the target sequence were identified using Benchling software. Three guide RNAs at each end of the target sequence were selected for activity testing. Guide RNAs were cloned into a T7 promoter vector followed by in vitro transcription and spin column purification. Functional testing was performed by transfecting a mouse embryonic fibroblast cell line with guide RNA and Cas9 protein. The guide RNA target site was amplified from transfected cells and analyzed by ICE (Synthego). One guide RNA at each end of the target sequence was selected, and a donor oligonucleotide was included to facilitate homologous recombination to produce a clean deletion event between the guide RNA cut sites. C57BL/6J zygotes were electroporated with 1.2 μM Cas9 protein, 47 ng/μl each guide RNA, and 400 ng/μl donor oligonucleotide and implanted in recipient pseudopregnant females. Resulting pups were screened by PCR and sequenced for the presence of the mutation allele. Male founders with the correct mutation were mated to wild-type C57BL/6J females for germline transmission of the mutated allele. Lines were back-crossed at least three generations.

### Histology and Immunohistochemistry

For histology and immunohistochemistry embryos were fixed in 4% paraformaldehyde/PBST overnight at 4°C. Embryos were then processed for either paraffin embedding or frozen in OCT. For histology, paraffin sections were dewaxed and stained with Hematoxylin and Eosin according to standard protocols. Histology sections were imaged on an Olympus BX61 fluorescence microscope. For immunohistochemistry, cryosections were thawed, washed in 1X PBS and subjected to antigen retrieval as previously described^62^. The following primary antibodies were used: mouse anti-tropomyosin (DSHB clone CH1), 1:50; rabbit anti-phospho-histone H3 (Millipore #06-570), 1:200; mouse anti-Islet1 (DSHB clone 39.4D5), 1:75; rabbit anti-Tle3 (ab94972), 1:500; rabbit anti smooth muscle actin (ab5694),1:500; rabbit anti Tbx20 (Genscript). Secondary antibodies were Alexa Fluor 488 goat anti-mouse IgG H+L (Thermo #A11001), 1:1000; Alexa Fluor 546 goat anti-mouse IgG1 (Thermo #A21123). Immunohistochemistry images were captured on a Zeiss LSM 700 laser scanning confocal microscope. Whole-mount images were captured using a Leica MZ 16F dissection microscope with a Retiga 4000RV camera. ImageJ (NIH) was used for image analysis and standard image processing.

### Scanning electron microscopy (SEM)

SEM was performed as previously described^62^. Briefly, embryos were fixed in paraformaldehyde/2.5% glutaraldehyde in 1X PBS, washed in 1X PBS, dehydrated and subjected to critical point drying. Embryos were mounted ventral side up and ion sputtered with gold palladium to 10mm thickness. Embryos were scanned with a Zeiss Supra 25 FESEM microscope. SEM photomicrographs were taken in standard orientations and magnifications.

### Quantitation of cardiomyocyte proliferation

For quantification of cardiomyocyte proliferation, embryos were processed as described above (see histology and immunohistochemistry methods). Cryosections were stained for TMY, pHH3 and DAPI. For quantitative analysis, 3-4 slides (4 sections per slide) were analyzed from wild-type (Tbx20^+/+^) (n=3) and Tbx20^EH1mut^ (Tbx20^EH1Mut/ EH1Mut^) (n=4) embryos. These sections represented the entirety of the heart (anterior, middle, and posterior). Mitotic index was calculated by dividing total cells positive for TMY and pHH3 by total cells positive for TMY. Statistical analysis was performed using student’s *t*-test.

### Quantitation of Islet1 positive cardiomyocytes and proliferating Islet1 positive cells in the second heart field

For quantification of Isl1-positive cardiomyocytes, embryos were processed as described above (see histology and immunohistochemistry methods). Cryosections were co-stained for Isl1, SMA and DAPI. A total of 7 sections corresponding to anterior, middle, and posterior positions of the heart were analyzed from wild-type (n=3) and Tbx20^EH1mut^ (n=3) embryos. Percent Isl1 positive cardiomyocytes was calculated by dividing total cells positive for SMA and Isl1 by total cells positive for SMA. To quantify mitotic index of Isl1 positive cells in the second heart field, cryosections were co-stained with Isl1, pHH3, and DAPI. A total of 7 sections corresponding to anterior, middle, and posterior positions of the SHF were analyzed from wild-type (n=4) and Tbx20^EH1mut^ (n=4) embryos. Mitotic index was calculated by dividing total SHF cells positive for Isl1 and pHH3 by total SHF cells positive for Isl1. Statistical analysis was performed using student’s *t*-test.

### RNA-sequencing and analysis

E9.5 hearts were collected from 12 WT and 12 Tbx20^EH1Mut^ embryos. 3 hearts were pooled per biological replicate to give a total of 4 Tbx20^+/+^ samples and Tbx20^eh1mut/eh1mut^ samples. RNA was isolated using RNAqueous micro kit (Ambion) as per the manufacturer’s protocol. Poly-A selected RNA-seq libraries preparation, sequencing reactions and initial bioinformatic analysis were conducted at GENEWIZ, LLC. Samples were run on a HiSeq2500 (Illumina) with 2×150 bp paired end reads.

Genes with an adjusted p-value < 0.05 and a log2(Fold Change) > 0.5 in either direction were considered differentially expressed. Canonical pathway and upstream regulator analysis was performed using Ingenuity Pathway Analysis (IPA) (QIAGEN Inc., https://www.qiagenbioinformatics.com/products/ingenuity-pathway-analysis). Tbx20 ChIP-sequencing data from E11.5 hearts was obtained from previously published data available in the European Molecular Biology Laboratory–European Bioinformatics Institute (EMBL-EBI) database (accession number E-MTAB-3967).

